# The high-resolution structure of cell membranes revealed by in situ cryo-electron tomography

**DOI:** 10.1101/2021.12.03.471052

**Authors:** Guanfang Zhao, Sihang Cheng, Yang Yu, Tianyi Zou, Huili Wang, Changlu Tao, Guoqiang Bi, Z. Hong Zhou, Hongda Wang

**Author notes:** Correspondence should be addressed to: G.-Q. B:; Or Z.H.Z; Or Hongda Wang. These authors contributed equally: Guanfang Zhao, Sihang Cheng, Yang Yu.

## Abstract

As the structural unit of life, cell is defined by the membrane system. The cell membrane separates the internal and external environment of the cell, and the endomembrane system defines the organelles to perform different functions^1-3^. However, lack of tools to *in situ* observe membrane proteins at a molecular resolution has limited our understanding of membrane organization and membrane protein interactions. Here we characterize the high-resolution 3D structure of human red blood cell (hRBC) membranes and the membrane proteins for the first time in situ by cryo-electron tomography (CryoET)^4-7^. By analyzing tomograms, we have obtained the first fine three-dimensional (3D) structure of hRBC membranes and found the asymmetrical distribution of membrane proteins on both sides of the membranes. We found that the membrane proteins are mainly located on the cytoplasmic side of hRBC membranes, with protein sizes ranging from 6nm to 8nm, in contrast to the ectoplasmic side with basically no proteins. Quantitative analysis of the density of hRBC membrane proteins shows that the membranes with higher protein occupancy have less phospholipid, making the membranes more rigid. Meanwhile, we obtained the channel protein-like structures by preliminary analysis of the membrane protein. Our results represent the first *in situ* structure characterization of the cell membranes and membrane proteins through cryoET and opens the door for understanding the biological functions of cell membranes in their physiological environments.

---

The complicated and highly organized molecular mechanism of cell membranes lays an important foundation for executing complex biological functions^8^. We imaged the prepared hRBC membranes by CryoET and the circular hRBC membranes with a diameter of 8-10μm can be identified clearly (Fig. 1a, b). The features of membranes together with other fine structure details become brighter with the 3D tomogram reconstructed (Fig 1c, d). Base on the research of hRBC membranes, we obtained the first overall morphology and structure of hRBC membranes through high resolution cryoET and observed the asymmetrical distribution of membrane proteins directly *in situ*. The gray scale on one side of the membranes are relatively homogeneous (Fig. 1e) when scanning through sections along the Z axis and almost no protein particles were observed (Fig. 1e1), indicating that the membranes on this side were smooth; however, a lot of membrane proteins were observed on the other side of membranes (Fig. 2f, f1), contributing to the roughness of membranes on this side, moreover, the residual spectrins on this side can also be located (Fig. 2g, g1). Combine the atomic force microscopy (AFM) data about hRBC membranes^9^, we concluded that the smooth side of the hRBC membranes was the ectoplasmic side and the other side coated with dense proteins was the cytoplasmic side of the hRBC membranes. In addition, by observing hRBC membranes from the XZ and YZ sections, we found that the length from one side to other side alone Z axis, spanning approximately 20 sections (Fig. 2h) or an estimated total thickness of about 10-15nm, consistent with the thickness of hRBC membranes^9^.

**Figure 1.**
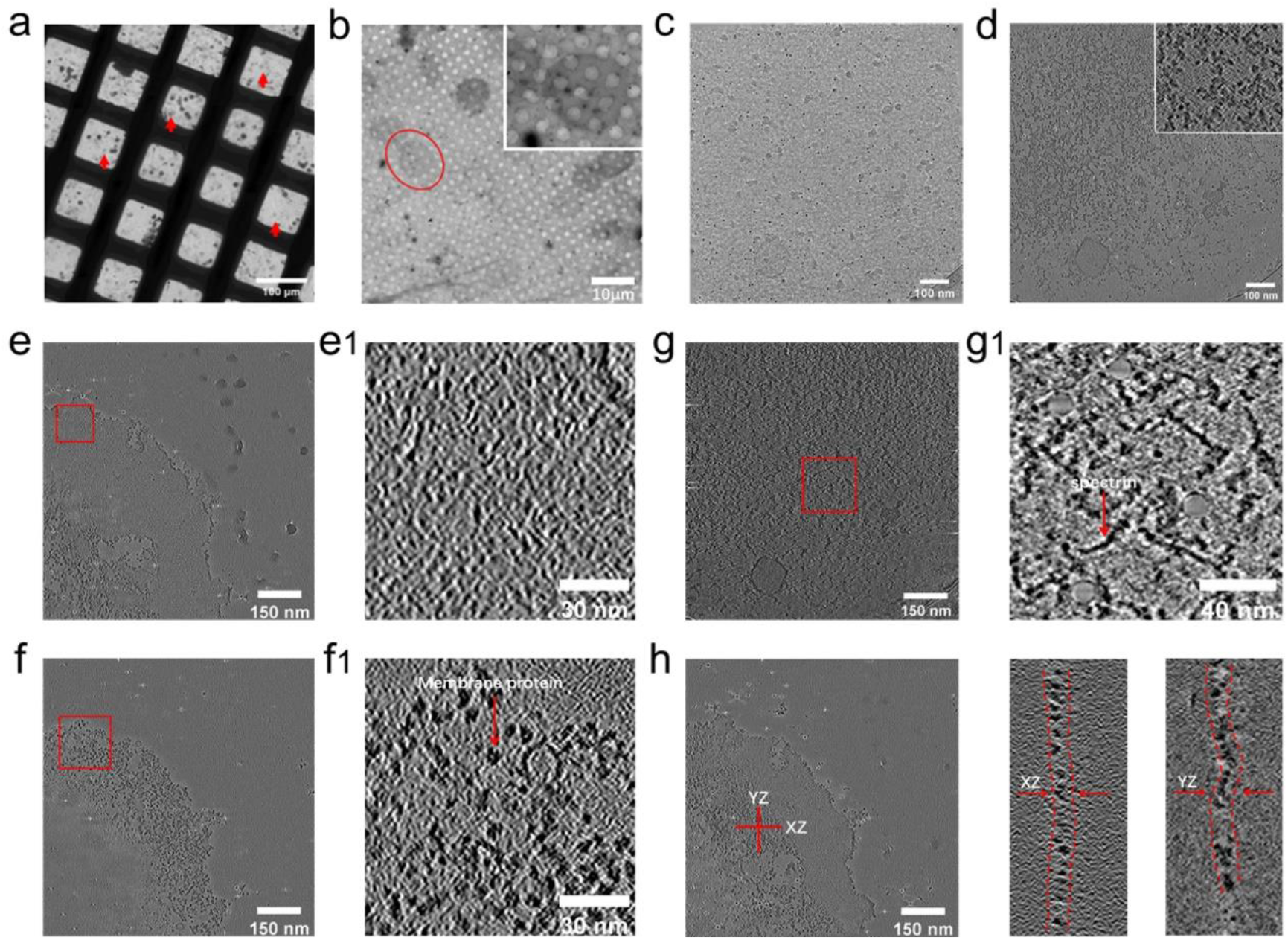
Overall morphology and structure of human red blood cell membranes. **a**, CryoET imaging of human red blood cell membranes at low magnification and the membrane patches can be recognized (red arrow). **b**, CryoET imaging in one grid square showing the membranes (red circle). Inset is the enlargement of membrane region marked by the red circle. **c**, A single cryoET tilt series projection image of membrane patch and nothing can be seen due to the low contrast. **d**, A slice from the 3D tomogram reconstructed from the tilt series of **c** showing structural details. Membrane proteins (red circle) can also be identified (inset image). **e, f**, CryoET images of the ectoplasmic and cytoplasmic side of the human red blood cell membrane respectively. **e1, f1**, Magnifying views of the red box area in B and C respectively show that the cytoplasmic side and ectoplasmic side of the human red blood cell membrane display completely different characteristics and many proteins are visible in the cytoplasmic side (red arrow in **f1**). **g**, The scattered distribution of spectrins on the cytoplasmic side of hRBC membranes. **g1**, Zoomed in view of the red box area in **g** and the spectrins can be visible obviously. **h**, Cutting surface of the hRBC membranes in the XZ and YZ direction.

Dense membrane proteins were coated on the intracellular hRBC membranes. We identified hundreds of tomograms and found that the distribution of membrane proteins on the cytoplasmic side of hRBC membranes had two different forms: sparse (Fig. 2a) and dense (Fig. 2b). With the membrane proteins identified in situ, we investigated their spatial distribution on cytoplasmic side of hRBC membranes. By measuring the distance between each membrane protein, we found that the nearest-neighbor (NN) distances of the membrane proteins in the sparse and dense areas are mainly distributed at 1.5-2.5nm and 0-1nm (Fig. 2c, d), respectively, suggesting that the membrane proteins are almost contact to each other in the dense areas. We then measured the concentration of membrane proteins, measured as the number of membrane proteins per μm^2^, in these two areas and suggested that the concentration of membrane proteins in the sparse areas was ranging from ∼4000μm^-2^ to ∼6000μm^-2^, while the concentration of membrane proteins in the dense areas was ranging from ∼10000μm^-2^ to ∼20000μm^-2^ (Fig. 2e, f). This also demonstrated the level of compactness of membrane proteins in the dense areas of hRBC membranes which provides basis for the biological functions of the cell membranes. Other than that, we randomly selected 100 sub-areas in the sparse and dense areas of cytoplasmic side of hRBC membranes, respectively, and followed by calculating the area of membrane proteins and phospholipids regions and found that the area percentage of membrane proteins was ranging from 20% to 30% in the sparse areas whereas it was ranging from 50% to 60% in the dense areas (Fig. 2g, h). This implied that the membranes had higher protein occupancy and less phospholipid content which would greatly weaken the fluidity of membranes^10^. This was in line with the reference’s statement “membranes are more mosaic than fluid”^11^. Beyond that, the membrane proteins in dense areas contacted closely with each other to form protein clusters and considered that the donut shape of human red blood cells caused cell membranes to endure a large membrane stress^12^, we believed that such dense proteins embedded in the inner membrane leaflets could, to some extent, serve as a local membrane skeleton to maintain the local membrane stress^13,14^. Intriguingly, when we randomly selected a sub-area from the dense areas (Fig. 2i) and labeled the position of membrane proteins precisely in that sub-area (Fig. 2j), we found that the membrane proteins tended to organize into clusters (the yellow circle in Fig.2i and 2j) and that clusters were spaced apart from each other (the white circle in Fig. 2i and 2j). Other techniques had illustrated that the membrane proteins tended to form functional microdomains (e.g. lipid raft) where proteins were densely arranged, consistent with the need for coordination to execute some biological functions^15-19^. However, further details about the membrane proteins in these areas were not available due to the resolution limit, but you were able to recognize individual membrane proteins in these areas of cryoET tomograms. At last, we plotted the length and width of all these membrane proteins, showing that it forms a cluster with the length chiefly distributed at 8nm and the width mainly distributed at 6nm (Fig. 2k).

**Figure 2.**
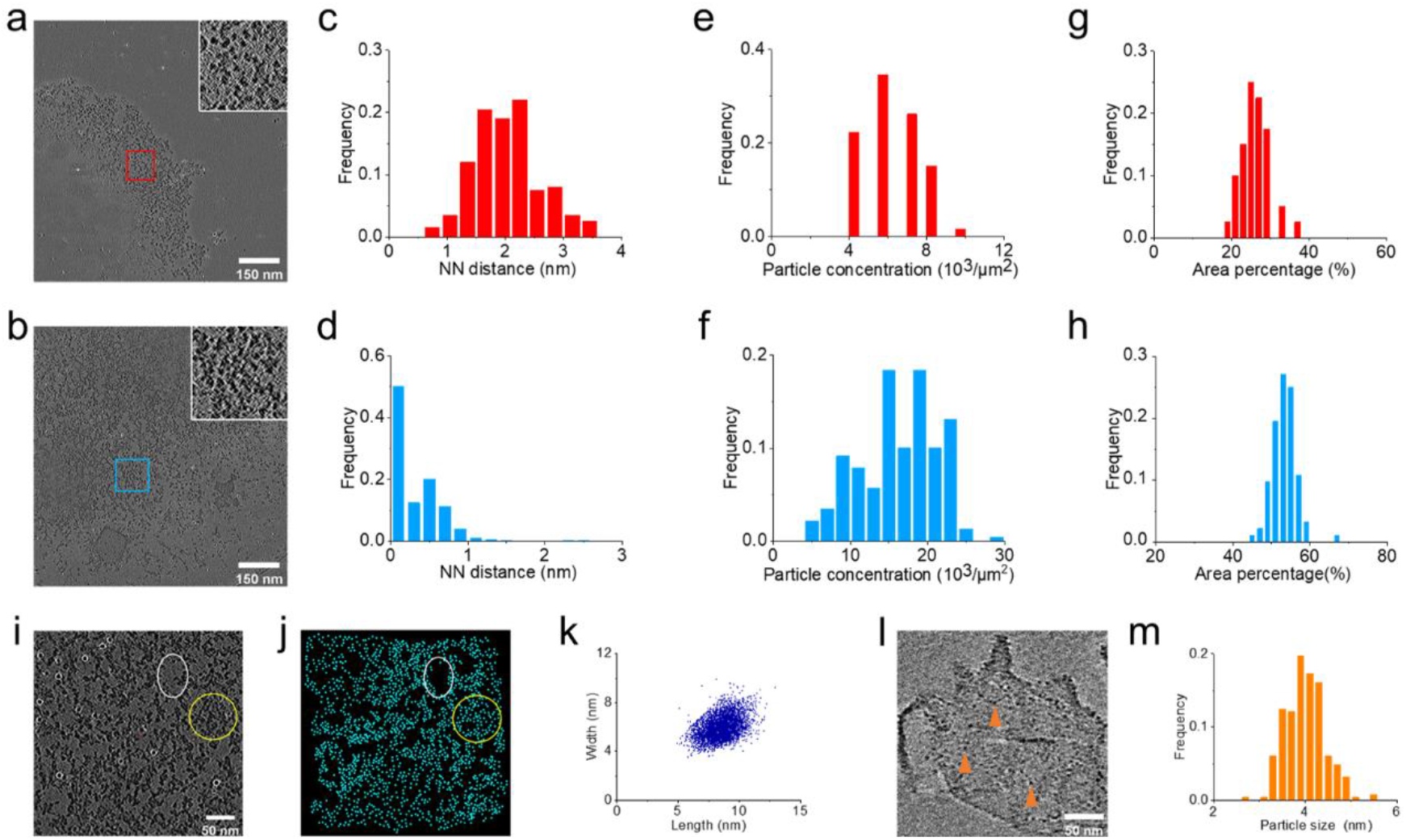
Characterization of membrane proteins on the cytoplasmic side of hRBC membranes. **a**, sparse areas and of membrane proteins in the cytoplasmic side of hRBC membranes. Inset is the zoom-in view of red box in **a** shows the sparse membrane proteins. **b**, dense areas of membrane proteins in the cytoplasmic side. Inset is the magnified view of light blue box in **b** shows the dense membrane proteins. **c, d**, The NN distance of membrane proteins in the sparse and dense areas. **e, f**, Concentration of membrane proteins in the sparse and dense areas. **g, h**, Area percentage of membrane proteins in the sparse and dense areas. **i, j**, A dense area of membrane proteins in the cytoplasmic side of hRBC membranes and the position of membrane proteins in this area. Membrane proteins formed clusters (yellow circle) with gaps between clusters (white circle). **k**, Length and width of the membrane proteins in the cytoplasmic side of the hRBC membranes. **l**, HRBC membranes digested by trypsin. The remained membrane proteins are labeled by orange arrows. **m**, Size distribution of remained membrane proteins after digesting the hRBC membranes by trypsin.

To further study the membrane proteins, we imaged the hRBC membranes after digestion. We found that some small granule densities were still remained in the membranes (Fig. 2l) that might be the separated peptides or transmembrane parts of membrane proteins, but the concentration of remained proteins was dramatically reduced compared to the untreated hRBC membranes. The size distribution of the remained proteins was in the range of 3-5nm with the peak at 4nm (Fig. 2m), which was also smaller than that of the untreated hRBC membranes.

Although the cell membranes and the membrane proteins can be identified clearly in the tomograms, the species of membrane proteins cannot be distinguished visually because of the small size of these membrane proteins and the effect of missing wedge. Many researches had shown that most of the proteins on the hRBC membrane are transmembrane proteins, with channel proteins (AE1, glut1, Na-K ATPase, etc.) making up the majority^20^. To identified the species of membrane proteins unbiased, we performed a template-free, Bayesian, three-dimensional (3D) classification with Relion for sub-tomograms selected by oversampling on the hRBC membranes (Fig. 3a, b). As shown in Fig 3c, we obtained a structure with a channel and this structure should be the channel protein in the hRBC membranes and this channel protein was ∼8nm in width and ∼9nm in length (Fig. 3d). we then extracted the sub-tomograms attributed to this channel protein and removing the duplicates, and a sub-tomogram average of in situ channel protein was obtained with a 25Å resolution through 3D refinement of Relion (Fig. 3e, f). The characteristics of the channel structure was particularly limited to the extent that it could not be distinguished to which channel proteins on the hRBC cell membranes it belonged at this resolution.

**Figure 3.**
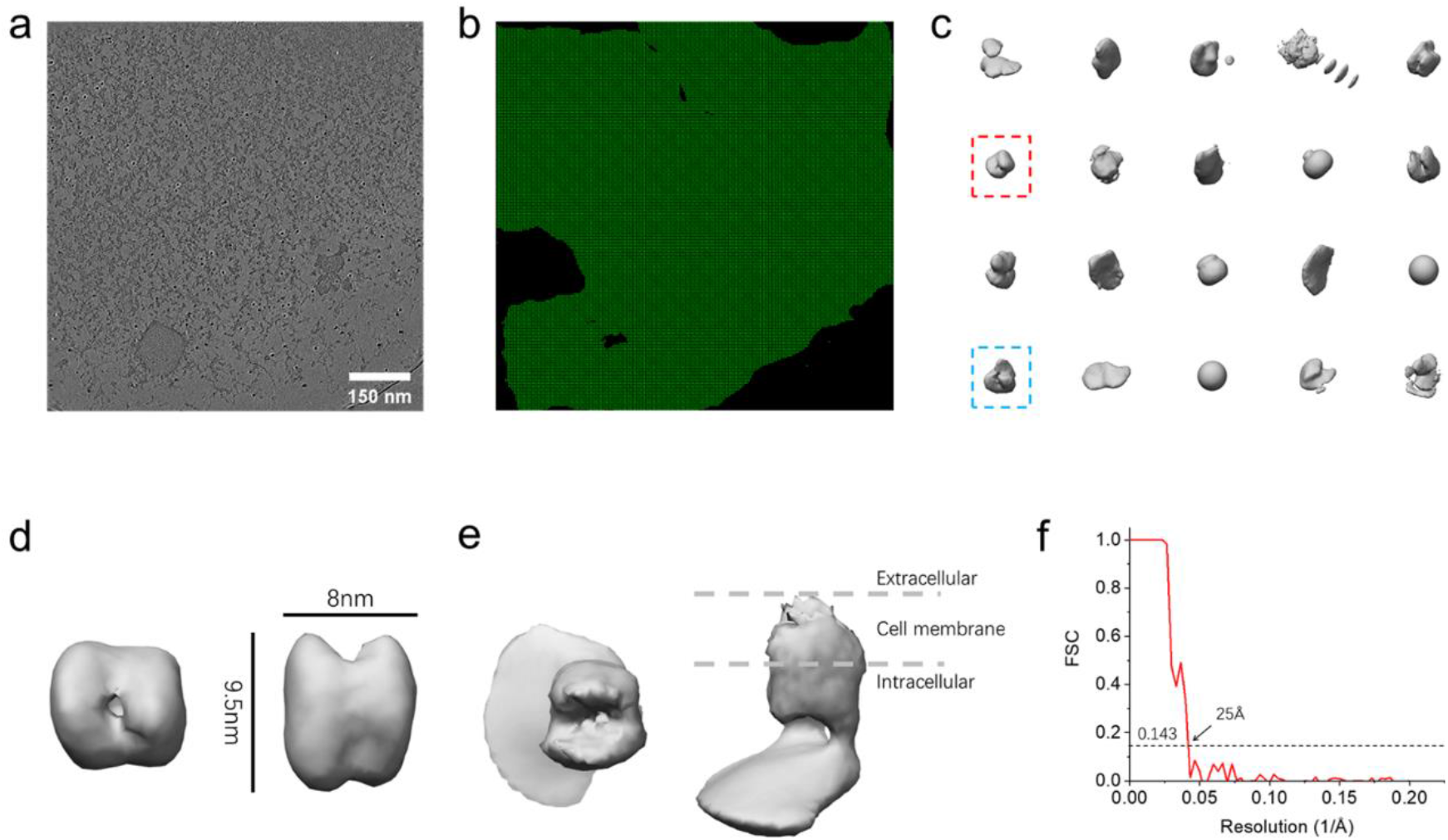
Identification of hRBC membrane proteins. **a**, Tomographic slice of an hRBC membranes. **b**, Sub-tomogram uniformly oversampling points on the hRBC membranes. **c**, Output structures of 3D classification. **d**, The channel structure in **c** (labeled by red box). **e**, Sub-tomogram average of the channel structure in **d. f**, Fourier shell correlation of the channel structure sub-tomogram average. FSC, Fourier shell correlation.

To analyze the spatial distribution of the channel protein of 3D refinement, we calculated the coordinate of the center of the sub-tomograms attribute to the channel protein and removed the duplicates, followed by plotting the distribution of coordinate points in the corresponding original tomograms (Fig. 4a, b). Subsequently, we used Ripley’s K function to quantitatively analyze the degree of clustering of these points^21^. The representative plot of average L(r) – r versus r shows a high level of clustering which suggests that the channel structures were organized into discrete nanoclusters on the hRBC membranes (Fig. 4c). Then we calculated the NN distance between these coordinate points and we found that the NN distance distributed at 8-15nm (Fig. 4d), suggested that these channel proteins were closely related to their neighbours if we considered the size of the channel proteins.

**Figure 4.**
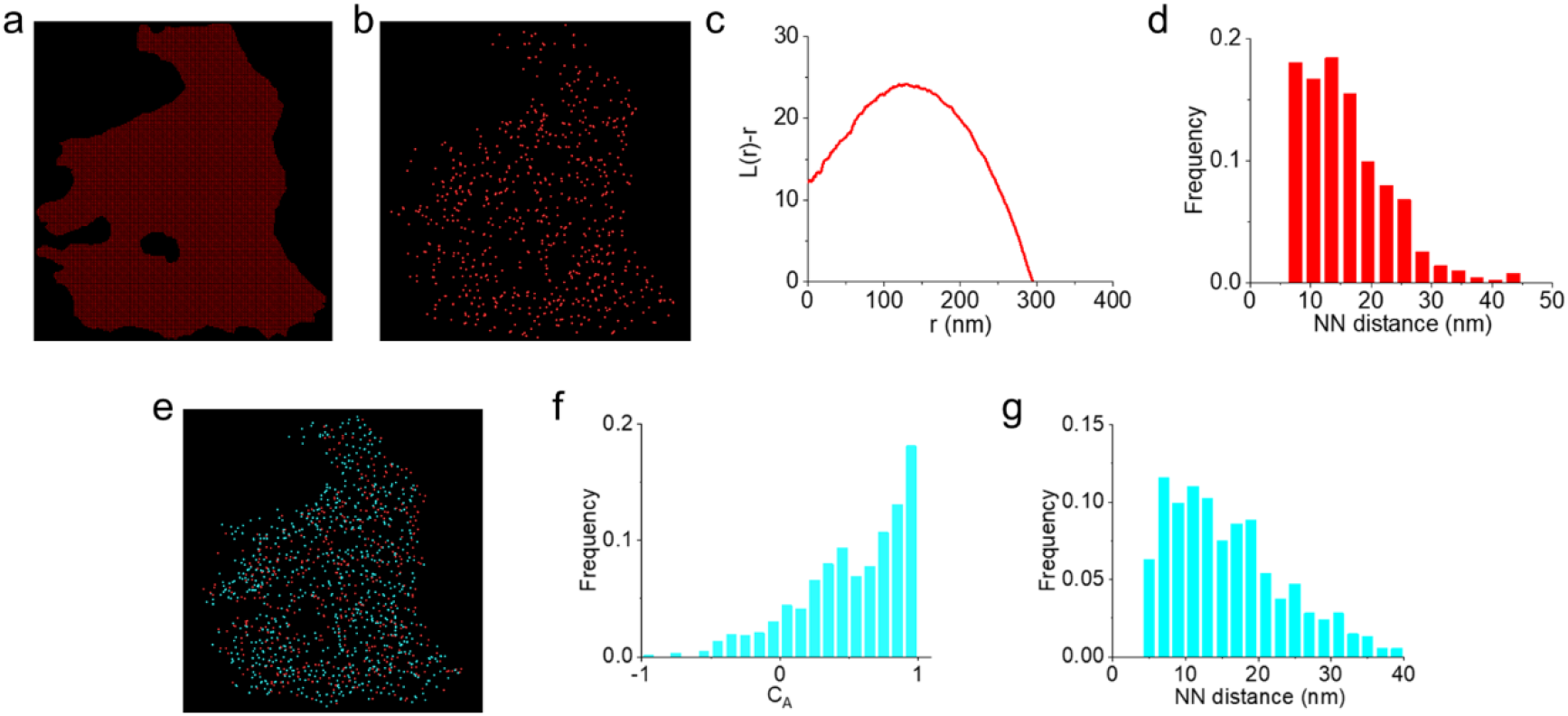
Quantitative analysis of spatial distribution of channel structure in hRBC membranes. **a**, Sub-tomogram uniformly oversampling points on the hRBC membranes. **b**, Distribution of the points belonging to the channel structure of 3D refinement. **c**, Ripley’s K function analysis of points belonging to the channel structure of 3D refinement. **d**, Distribution of the NN distance between the points belonging to the channel structure of 3D refinement. **e**, Merged image of the distribution of points belonging to the channel structure and points of another channel structure (labeled by light blue box). **f**, Distribution of the colocalization value C_A_ for the points of these two channel structures. **g**, Distribution of the NN distance between these two channel structures.

In addition, we extracted the coordinate points of sub-tomograms belonging to another channel structure (labeled by light blue box in Fig. 3c) with lower abundance in the output result of 3D classification and eliminated duplicates. We first merged the distributions of these two channel proteins in the same tomogram (Fig. 4e) and found that these points were distributed around each other. We then quantified the degree of spatial association between the two channel proteins by performing colocalization analysis using coordinate-based-colocalization (CBC) method^22^. It could be found that the colocalization value C_A_ for the two channel proteins revealed a maxima near 1, implying a strong colocalization between the two channel proteins (Fig. 4f). Finally we calculated the NN distance between the two channel proteins and found that it was mainly distributed at 8-13nm.(Fig. 4g). In short, the channel proteins tended to form protein clusters in the hRBC membranes and contacted closely with other channel proteins. This fitted well with the distribution characteristics of channel proteins in the hRBC membranes, which tended to interact with other proteins and form functional microdomains to perform biological functions^18^.

## Methods summary

The CryoET micrographs of frozen-hydrate hRBC membranes was collected from -60 to +60° at 3° intervals by Tecnai F20 transmission electron microscope (FEI) equipped with K2 Summit direct electron detector (K2 camera, Gatan). The acceleration voltage of Tecnai F20 was 200 KV. and the total electron dosage of ∼140 e^-^/Å^2^. The final pixel size is 0.2508 nm. Three-dimensional tomograms were reconstructed using IMOD^23,24^. The template-free, Bayesian, three-dimensional (3D) classification and 3D refinement was performed by Relion^25-27^. To analyzing the distribution of the channel proteins after 3D refinement. The Ripley’s K function was used to perform the cluster analysis of sub-tomograms belonging to channel proteins of 3D refinement^21^. To quantify the degree of colocalization between sub-tomograms belonging to different channel proteins, the Coordinate-based Colocalization (CBC) analysis was employed and calculate the CBC value (C_A_) which indicates a colocalization for 0 < C_A_ <1, no colocalization for C_A_ = 0 and segregated but near localization for -1 < C_A_ <0^22^. The NN distance was calculated for the membrane proteins using the standard distance formula in 3D.

## Acknowledgments

This work was financially supported by the National Key R&D Program of China (2017YFA0505300), National Natural Science Foundation of China (21727816, 21721003 and 21773225), Laboratory for Marine Biology and Biotechnology, Pilot National Laboratory for Marine Science and Technology (Qingdao) (MS2018NO08).

## Author Contributions

G.Z. S.C. Y.Y. performed the experiments. S.C. and Y.Y collected the data. G.Z. processed and analyzed the data and wrote the manuscript. T.Z. HL.W. participated in data processing. C.T assisted with data collection and results discussion. G.B. Z.Z. and H.W. conceived and designed the experiments and discussed the results and commented on the manuscript.

## Author information

The authors declare no conflicts of interest.

## Notes

### Competing Interest Statement

The authors have declared no competing interest.

